# A systematic review and ALE meta-analysis of cognitive control, motivation and effort-based decision-making in schizophrenia and mood disorders: Implications for multidimensional apathy

**DOI:** 10.1101/2025.01.23.634295

**Authors:** Giulia Lafond-Brina, Ludovic C. Dormegny-Jeanjean, Anne Bonnefond

## Abstract

**Background:** Apathy is present in about 50% of patients suffering from schizophrenia (SZ) and mood disorders (MD). Even though these two disorders are different, in psychiatry, apathy refers typically to the same unidimensional clinical entity, sharing the same pathophysiological processes and symptoms regardless of the underlying diagnosis. Neurology proposes another perspective: three forms of apathy —emotional, executive, and initiative— are related to a disturbance in motivational processing, cognitive control, and effort-based decision-making, respectively. We explored whether this latter model can be applied in psychiatry by identifying differences between SZ and MD through a PRISMA meta-analysis of imaging studies on motivational, cognitive control, and effort-based decision-making networks.

**Methods:** We searched the Sleuth BrainMap database for studies on SZ, MD, and/or healthy controls using tasks that explore motivation, cognitive control, and effort-based decision-making. Twenty-eight functional MRI studies were identified and included for a coordinate-based activation likelihood estimation analysis.

**Results:** In SZ, hypoactivity in the motivational structures, a specific hyperactivity in the right cerebellar vermis that has previously been linked to emotional blunting and apathy, and a lack of activation during effort-based decision-making that could imply an impaired reward processing, suggest a dominant form of emotional apathy. In MD, hypoactivity in both cognitive control and motivational structures, which has previously been linked to the co-occurrence of executive difficulty and amotivation, suggests a dominant mix of emotional/executive apathy.

**Conclusions:** Despite a small number of studies, our results could help target new individualized treatment strategies in precision psychiatry based on a multidimensional approach to apathy.

**Graphical abstract:** 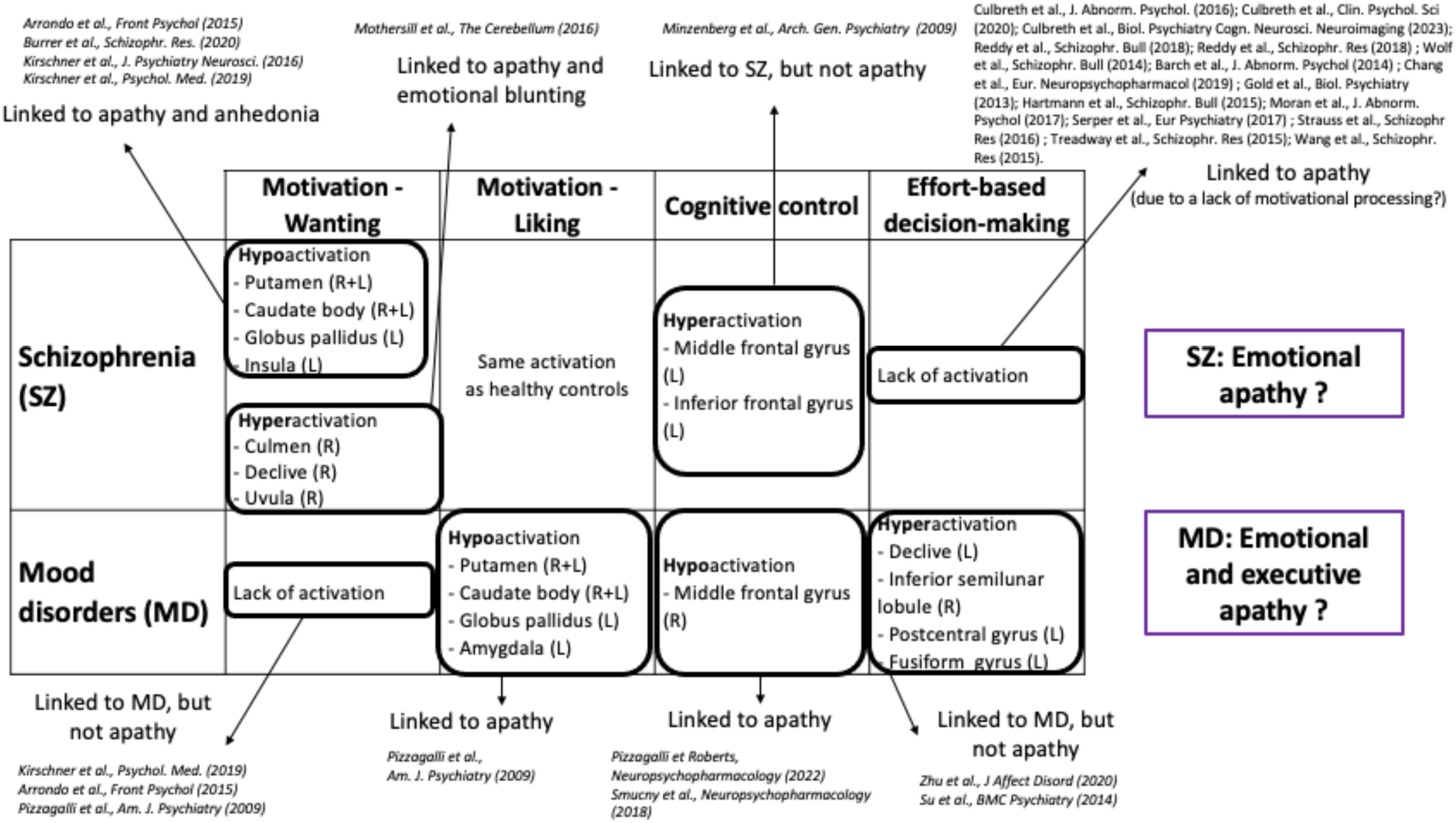

The results of this meta-analysis highlight different activation profiles in SZ and in MD during motivational, cognitive control, and effort-based decision-making tasks, suggesting a dominant form of emotional apathy in SZ and a dominant coexistence of emotional and executive apathy in MD.

## Introduction

Apathy, a behavioral symptom that often appears during the prodromal phase of psychiatric disorders, is prevalent in patients suffering from both psychotic disorders (including schizophrenia [SZ]) and mood disorders [MD] [1–5]. Paradoxically, apathy, which is extremely burdensome for these patients, has been poorly studied in psychiatry, and dedicated treatment options remain scarce [6,7].

Why does psychiatry need to reconsider its common approach to apathy? Historically, psychiatry of the early 20^th^ century described a precise semiology of apathy and motivational disorders with a multitude of symptom-specific terms: “avolition” for lack of initiative, “apragmatism” for inability to conduct daily basis actions, and “athymhormie*”* (i.e. the loss of psychic self-activation conceptualized by the French psychiatrists Dide and Guiraud) (see a review in [8]). This detailed semiology carried certain hypotheses, including that the various apathetic symptoms may be supported by specific pathophysiological processes that may interact with each other. By extension, these symptoms and the pathophysiological processes associated with them may be transnosographic for some, and specific to MD or SZ for others (e.g., according to Delay’s concept of “hypothymie*”*, SZ patients seem indifferent to their emotional blunting, whereas some MD patients have been described as paradoxically uncomfortable with their emotional blunting, which can be the basis of their complaint, as in the apathetic depression of the Wernicke-Kleist-Leonhard classification [9,10]).

However, the atheoretical conception of modern psychiatric classifications (i.e., the DSM and the ICD) has replaced these systems-based neurology approaches [11]. In this context, semiological terms used for apathy-related symptoms in various disorders are now considered almost synonymous and even sometimes used without distinctions, while, paradoxically, the possibility that they may involve transdiagnostic pathophysiological pathways or equifinality has been bypassed (see a review in [12]). As apathy is still underdiagnosed and underrecognized as the outcome of specific pathophysiological processes, psychiatry lacks specific therapeutic strategies [6]. Some treatments may even worsen these processes (e.g. selective monoamine reuptake inhibitor in MD [7,13,14]), which can result in apathy as a chronic residual symptom. Latest neuroscience-based approaches (e.g. the Research Domain Criteria [RDoCs] initiative) propose modeling the link between pathology, behavior, and neurobiology using transdiagnostic constructs (see a review in [15]). However, the RDoCs initiative is still seeking a model for practical application in the clinical setting.

Levy and Dubois’s (2006) neurological model of apathy may be a good compromise to overcome these limits while retaining operationalized and practical model. Indeed, based on clinical observations of neurological patients with brain lesions affecting the prefrontal cortex (PFC) and the basal ganglia, three forms of apathy —emotional, executive, initiative— related to a disruption of “emotional–affective”, “cognitive” or “auto-activation” processing, respectively, have been identified [16–18]. As such, the existence of a dominant form of apathy is assumed to be related to a neurological system dysfunction, irrespective of the overall diagnosis. Indeed, emotional apathy, i.e. an emotional blunting and indifference to emotional and motivational information, reflects a specific impairment of the motivational network (orbital and ventromedial PFC, as well as the limbic territories of the basal ganglia, especially the ventral striatum; [19–21]). Executive apathy, i.e. a difficulty to plan an action, suggests an impairment of the cognitive control network (dorso/ventrolateral PFC), as well as the cognitive territory of the basal ganglia (especially the dorsal caudate nucleus and the internal portion of the globus pallidus); [22–24]). Initiative apathy, i.e. a difficulty to initiative thought, action or emotion impacting effort-based decision-making [25], suggests an impairment of both motivational and cognitive control networks, with specific alterations in the anterior cingulate cortex and the supplementary motor area [26–31].

Studies in SZ and MD have already explored the potential links between functional impairments of cognitive processes and apathy without a physiopathological hypothesis. In this sense, concerning motivational processing, a body of evidence exists in favor of preserving ability to experience immediate pleasure (i.e. liking related processes) in SZ, but impaired ability to hedonic processes that involve a time delay (e.g. in wanting related processes) (see a review in [32]). However, more mixed evidence exists in MD, with clinical studies pointing toward impaired wanting-related processes [33], whereas fMRI studies identify both wanting and liking deficits [15,34,35]. Concerning cognitive control processing in SZ, several studies have shown an inability to anticipate and actively represent goal information needed to guide behavior (i.e. the deficit of proactive cognitive control process; see a review in [36]), whereas in MD, the evidence is more heterogeneous and points to cognitive control deficits that can be more proactive or reactive depending on the context and the emotional and motivational state of the subjects [37–39]. Concerning effort-based decision-making processing, in SZ, evidence suggests an altered processing, with a reduced (physical or cognitive) effort expenditure only for high reward [40–51,51–63]. In MD, several studies showed an impairment of (physical or cognitive) effort-based decision-making processing, with a higher subjective effort cost regardless of the reward [62–69]. However, one study found the same decision-making choices in MD as in healthy subjects [61].

Highlighting the dominant form of apathy in SZ and MD could be a first step to providing some more precise hypotheses in terms of the impaired mechanisms behind apathy in these two disorders. If mechanisms underlying emotional, executive, and initiative apathy are transdiagnostic, a first argument for this hypothesis should be a cross-diagnostic continuum between SZ and MD, with impaired (dis)liking network for emotional apathy, altered cognitive control network in executive apathy, and impaired effort-based decision-making network in initiative apathy regardless of the pathology. On the contrary, there will be a first argument in favor of pathology-specific mechanisms of apathy if distinct motivational, cognitive control, and decision-making networks are highlighted in SZ and MD.

To do this, the development of meta-analysis methods for neuroimaging data provides a valuable tool for combining data across diverse studies and building consensus in the identification of neuroanatomical correlates of specific processes. Activation likelihood estimation (ALE) metanalysis, one of the most promising and reliable statistical meta-analysis approaches, weights the foci by the number of participants in each study, identifies common activations across different studies, and uses random-effects inference [70–72]. In order to limit the risk of bias in the results, by showing different types of perceptual and cognitive processes due to the use of several task paradigms [73], this meta-analysis will only focus on one motivational task and one cognitive control task: the Monetary Incentive Delay (MID) task [74] and the Dot Probe Expectancy version of the AX-Continuous Performance Test (AX-CPT) task [75] respectively. Both tasks have already been confirmed to probe inter-individual and cross-diagnostic differences in psychiatric disorders [76,77]. Moreover, by using these two tasks coupled with electroencephalography, we recently highlighted, with healthy subjects presenting either an emotional or an executive apathy phenotype, specific impairments to each form of apathy. A disliking impairment was associated with emotional apathy and a proactive cognitive control deficit with executive apathy [78]. Nevertheless, regarding decision-making, no consensus-based task or paradigm exists to assess (physical or cognitive) effort-based decision making [79–81]. For that reason, any task based on the assessment of the amount of effort a person is willing to exert to obtain a certain level of reward will be accepted in this meta-analysis.

Therefore, an ALE meta-analysis of neuroimaging studies of motivational, cognitive control, and effort-based decision-making processes in SZ and MD during the completion of tasks, discussed in regard to the known neural substrates of multidimensional apathy, thus seems like a promising approach to investigating the existence of dominant forms of apathy in psychiatric disorders. The objective of the present meta-analysis is to determine the extent to which the observed functional activations of motivational, cognitive control, and effort-based decision-making networks in SZ and MD could suggest the existence of dominant forms of apathy specific to each of these psychiatric disorders.

## Methods and materials

### Literature selection

A search of the Sleuth BrainMap database [72] was performed to identify all English-language, peer-reviewed studies that reported fMRI activations related to motivation or cognitive control or effort-based decision-making in healthy controls (HC) and patients suffering from SZ and/or MD, using fMRI or positron emission tomography (PET). The online research was conducted using the diagnosis keywords ‘Schizophrenia’ or ‘Major depressive disorder’ or ‘Bipolar disorder’ or ‘Healthy controls’ and the paradigm keywords ‘Reward’ or ‘Go/no go’ or ‘n-back’ or ‘button press’. No other research filter was used. In addition, all the references of retrieved studies and pertinent review articles were manually checked.

To investigate motivational processes, we only examined studies that used the MID task. The MID task is a simple detection task that allows the exploration of the two motivational components, the wanting, *i.e.* the anticipation of a reward, and liking, *i.e.* the feeling of pleasure when obtaining that reward [74,82,83] (Figure 1a). During the MID, the wanting is investigated by the reward or loss cue vs neutral cue during the anticipation phase; the liking during the receipt phase.

**Figure 1:**
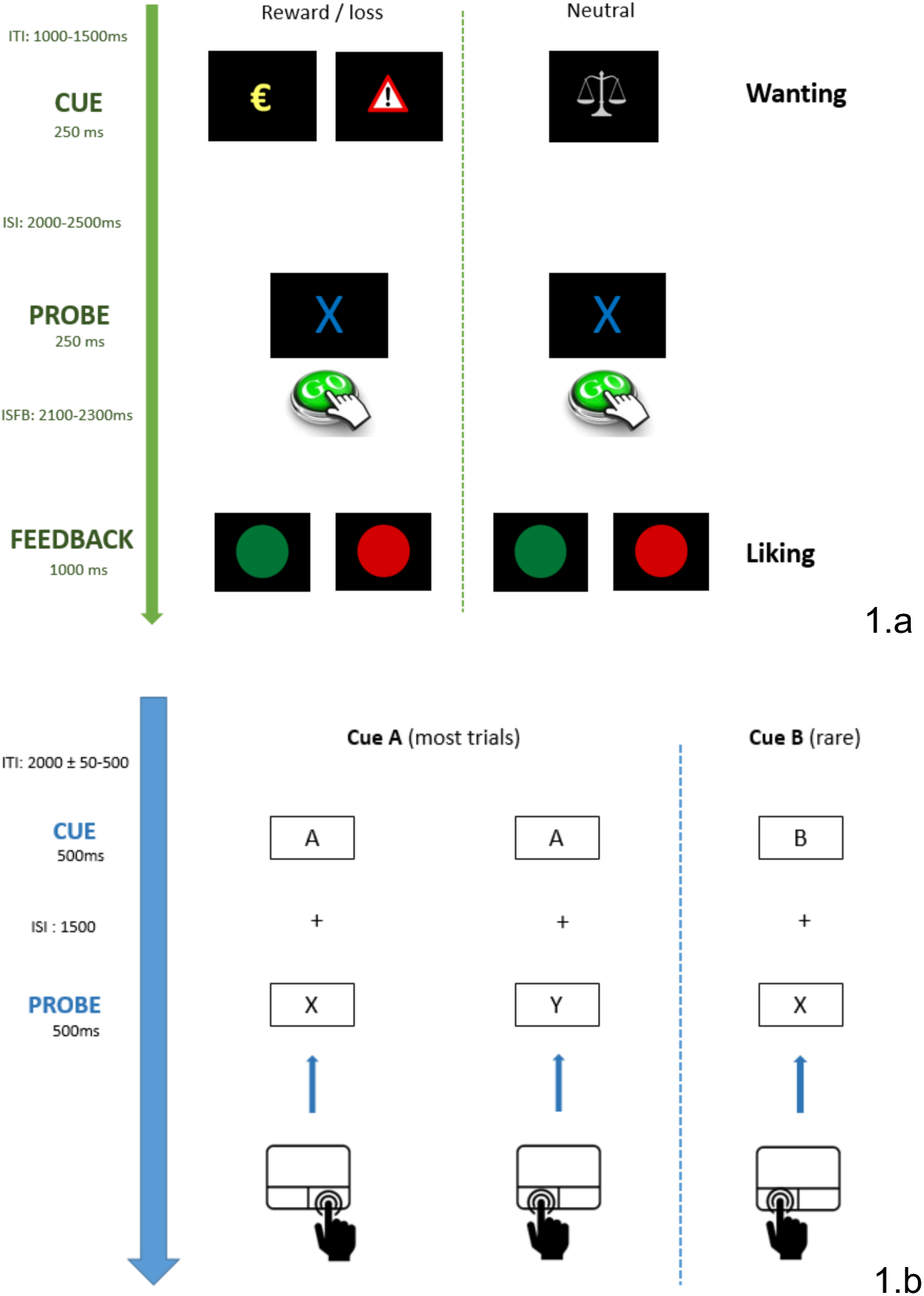
The MID and the AX-CPT tasks. 1. a: The MID is a visual detection task in which subjects are required to respond as quickly as possible by pressing a response button at the presentation of the probe (a cross generally). An incentive (reward or loss) or neutral cue (no reward and no loss) is presented before the probe. Either positive or negative (a green or red disc generally) feedback (FB) is given after the response. The FB informs the participant if his or her response was completed with enough speed, and in case of incentive cues about the obtained reward or loss. After a reward cue, positive FB indicates an actual monetary reward, whereas negative FB indicates the absence of a monetary reward. After a loss cue, positive FB indicates the absence of monetary loss, whereas negative FB indicates an actual monetary loss. After a neutral cue, positive and negative FB indicates no monetary variation. 1. b: The AX-CPT is a cue-probe detection task in which letters are presented one by one on the screen: A or B cues alternated with combinations of X or Y probes. The subjects had to respond after the probes by pressing the right button in cases presenting the target sequence (“A cue – X probe” trials). In all other cases, they had to press the left button. Thus, four types of trials were differentiated: AX, AY (where Y is any other combination apart from X), BX (where B is any other combination apart from A), and BY trials. The target sequence (AX) was more frequent (about 70% of trials) than the other sequences (AY, BX and BY), leading participants to develop a strong expectation of a ‘X’ probe following a ‘A’ cue.

To investigate cognitive control, we only examined studies that used the AX-Continuous Performance Test (AX-CPT) task. The AX-CPT is a cue-probe detection task that is the most used paradigm to explore proactive and reactive cognitive control processes [75], corresponding respectively to a control mode engaged before *versus* at the moment of target appearance [84,85] (Figure 1b). During the AX-CPT, the A vs B cues reflect the proactive cognitive control, whereas the reactive cognitive control is based on AX vs AY vs BX trials.

For the current meta-analysis, the following inclusion criteria for fMRI studies were utilized:

1. subjects were healthy controls or patients diagnosed with either MD or SZ;
2. MD or SZ were diagnosed according to DSM-III, DSM-IV(−TR) or DSM-5;
3. studies reporting imaging results of activations and deactivations using blood oxygen level-dependent (BOLD) fMRI (1.5 or 3T) with whole brain analysis;
4. studies using the MID (wanting and liking) and/or the CPT-AX (cognitive control) and/or an effort-based decision-making task as experimental fMRI paradigms and reporting first-level analysis results (i.e. paradigm-related within group contrasts);
5. studies contrasted directly two active conditions (e.g., monetary gain or loss vs neutral cue, B vs A cues, high effort vs low effort);
6. coordinates were reported in either standard Talairach space or Montreal Neurologic Institute (MNI) space.

Studies were excluded if:

1. the subject pool overlapped with other published studies on smaller subsets of the same sample (in that case, the subject included was the one with the larger sample size).
2. there was no whole-brain analyses;
3. coordinates could not be retrieved;
4. comparisons included a resting state condition.

No age, gender or treatment restrictions were applied.

### ALE meta-analysis procedure

Coordinates from motivation and cognitive control studies were analyzed separately following the Activation Likelihood Estimation (ALE) technique implemented in GingerALE 3.0.2 (http://brainmap.org/ale/, RRID: SCR_014921). This version uses a random effect model and weighting for sample size of the original studies [70,86]. Coordinates of the foci of activation reported in the original studies were transformed into Talairach space using the Lancaster transform (icbm2tal tool) in GingerALE [87]. In ALE, activation foci reported in original studies are treated as 3D Gaussian distributions centered at the reported coordinates. Activation probabilities are then calculated for each standard-space voxel to construct ALE maps for contrasts of interest. To determine the reliability of the ALE map, null-distributions are generated by analyzing the distribution of ALE values across independent studies, which is conceptually similar to using permutation tests of individual voxels across experiments. The observed values in the ALE distribution are then compared to the null distribution in order to assign probability estimates to the experimental data.

Three ALE analysis sets, based on paradigm related within-group contrasts maps for each group (MD, SZ and HC) were realized. Then, for comparisons, the ALE maps for SZ and MD for each experimental design were contrasted using subtraction meta-analysis procedure implemented in GingerALE [86]. For all analyses, only clusters exceeding the p = 0.05 FWE-corrected significance threshold were considered.

For visualization, whole-brain maps of ALE maps were imported into multi-image analysis GUI (MANGO; http://ric.uthscsa.edu/mango, RRID: SCR_009603) and overlaid onto a standardized anatomical template in Talairach space (Colin 1×1×1, available in www.brainmap.org/ale).

## Results

1623 records were identified in the Sleuth database and 28 studies were included in the meta-analysis (see the flow diagram of studies selection process in Figure 2).

**Figure 2:**
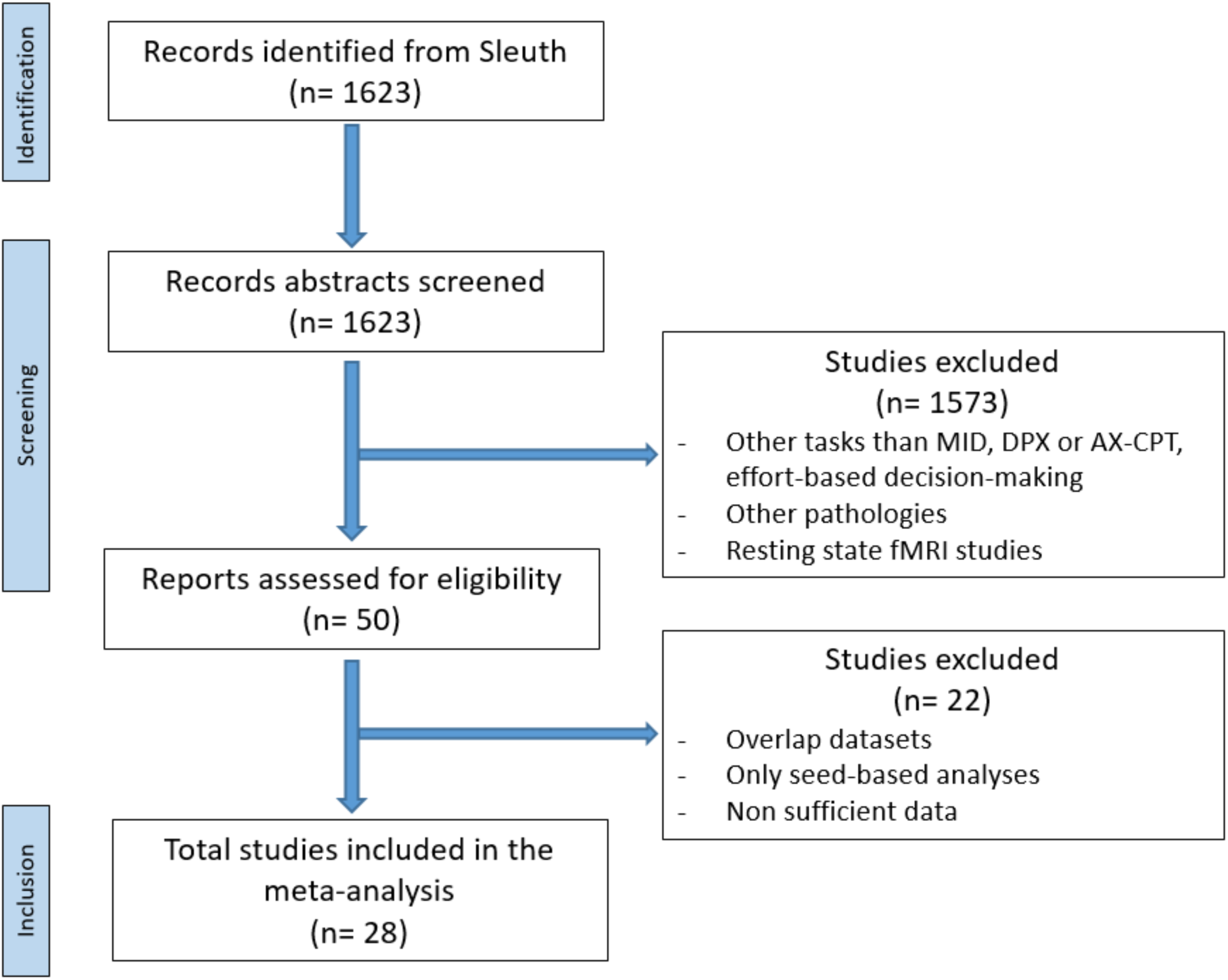
PRISMA flow diagram of the identification of articles.

Identification details of included studies, within each domain, are reported in Table 1a. The sample characteristics for each study are summarized in Table 1b.

**Table 1a:**
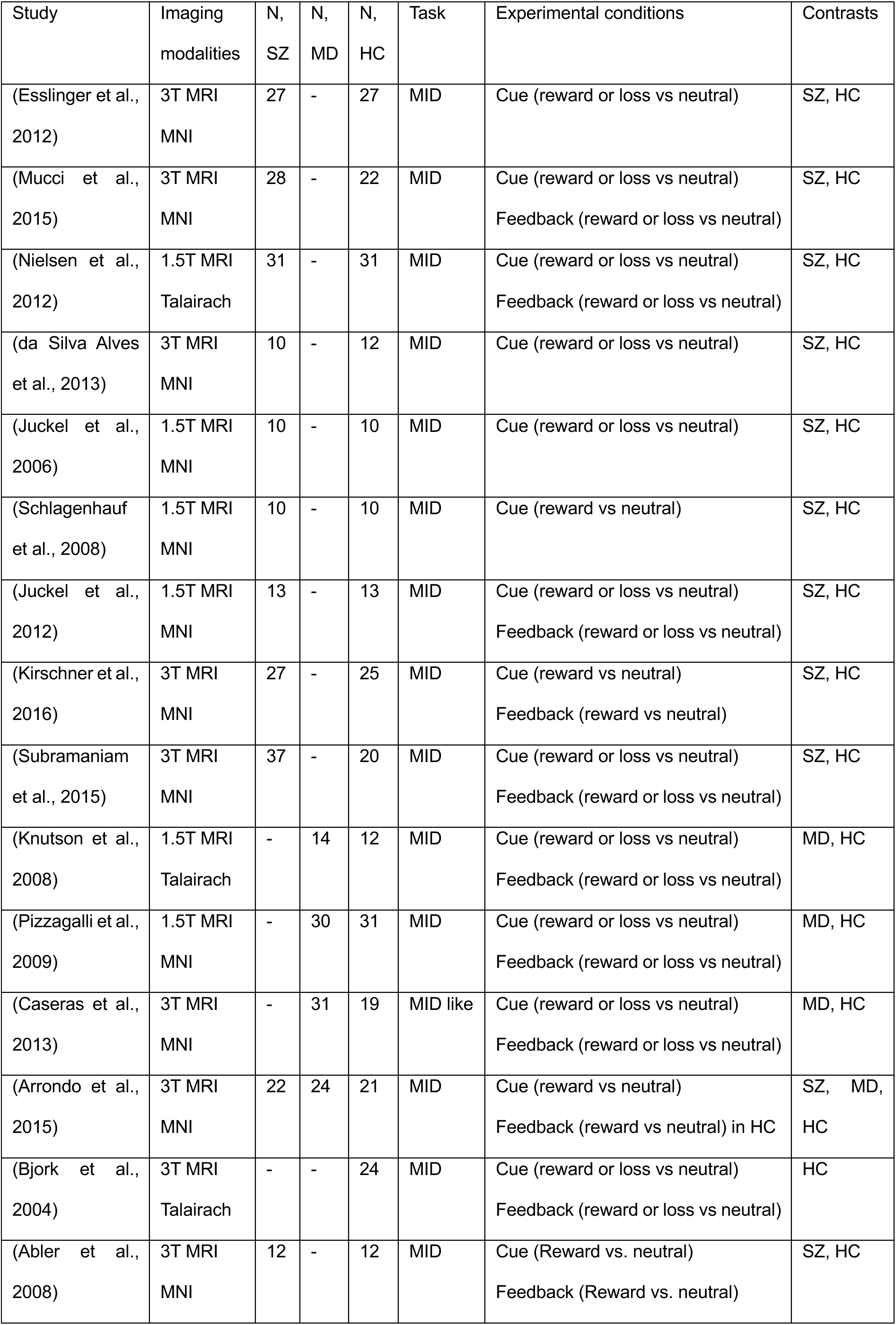
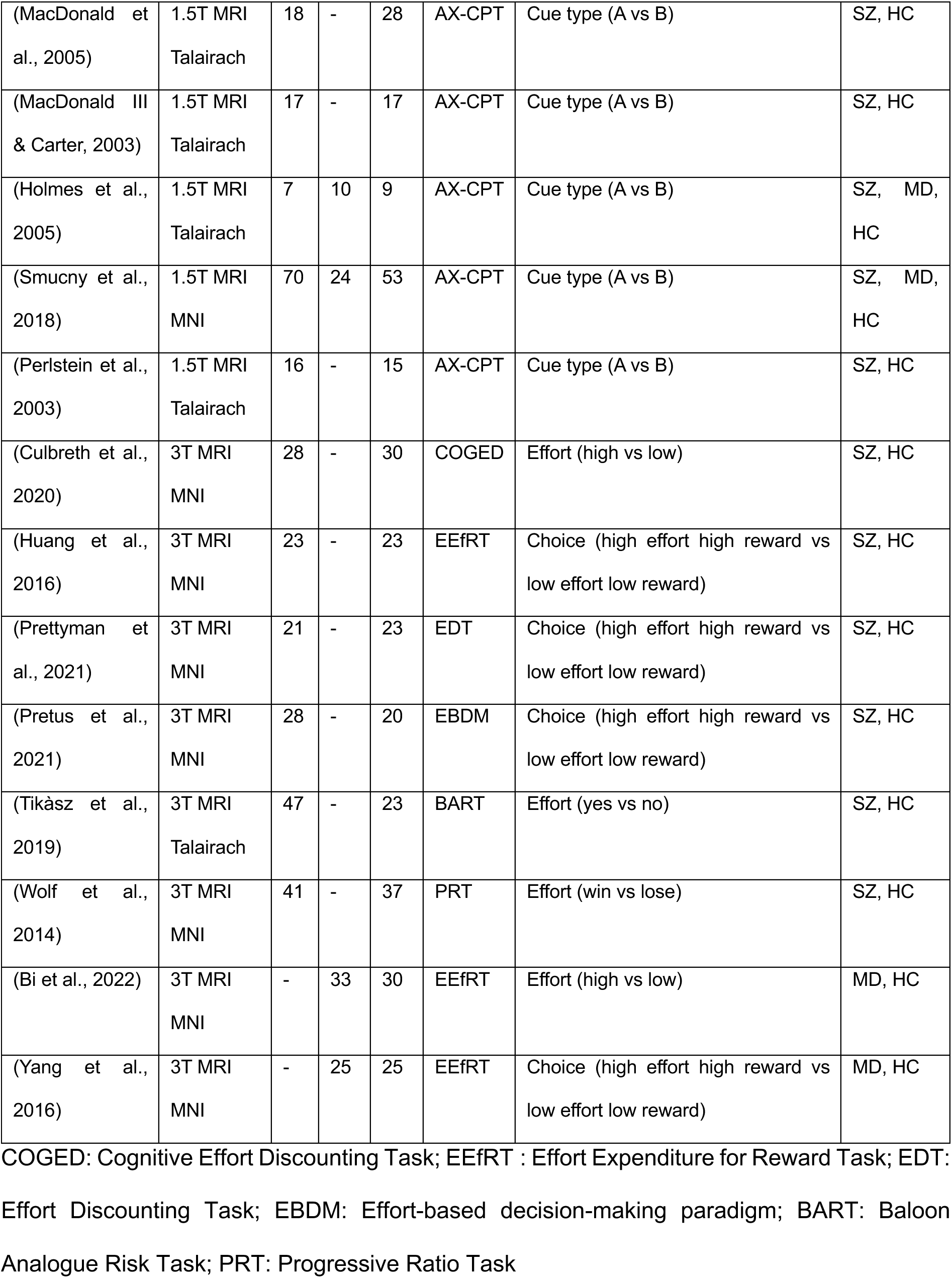
Study information for all papers included in the meta-analysis.

**Table 1b:** Participants information for all papers included in the meta-analysis.

| Study | N |  |  | Age |  |  | Gender (M/F) |  |  | Treatments<br>(mg of CLPZ<br>equivalents) |  |
| --- | --- | --- | --- | --- | --- | --- | --- | --- | --- | --- | --- |
|  | SZ | MD | HC | SZ | MD | HC | SZ | MD | HC | SZ | MD |
| (Esslinger et al., 2012) | 27 | - | 27 | 27.8<br>(7.4) | - | 27.1<br>(5.9) | 20/7 | - | 20/7 | 0 | - |
| (Mucci et al., 2015) | 28 | - | 22 | 33.1<br>(6.7) | - | 31.9<br>(8.5) | 18/10 | - | 10/12 | ? | - |
| (Nielsen et al., 2012) | 31 | - | 31 | 25.9<br>(6.4) | - | 25.7<br>(5.8) | 22/9 | - | 22/9 | 0 | - |
| (da Silva Alves et al., 2013) | 10 | - | 12 | 22.7<br>(3.2) | - | 34.6<br>(11.2) | 10/0 | - | 12/0 | ? | - |
| (Juckel et al., 2006) | 10 | - | 10 | 26.8<br>(7.8) | - | 31.7<br>(8.4) | 10/ 0 | - | 10/ 0 | 0 | - |
| (Schlagenhauf et al., 2008) | 10 | - | 10 | 30.5<br>(10.6) | - | 31.8<br>(8.7) | 9/1 | - | 9/1 | 189 (8) | - |
| (Juckel et al., 2012) | 13 | - | 13 | 25.5<br>(4.6) | - | 25.7<br>(4.8) | 11/2 | - | 11/2 | ? | - |
| (Kirschner et al., 2016) | 27 | - | 25 | 31.9<br>(7.1) | - | 33.0<br>(9.7) | 18/9 | - | 16/9 | 491<br>(349) | - |
| (Subramaniam et al., 2015) | 37 | - | 20 | 45.1<br>(9.9) | - | 43.7<br>(13.3) | 25/12 | - | 14/6 | 375<br>(555) | - |
| (Knutson et al., 2008) | - | 14 | 12 | - | 30.7<br>(8.8) | 28.7<br>(4.3) | - | 5/11 | 4/8 | - | 0 |
| (Pizzagalli et al., 2009) | - | 30 | 31 | - | 43.2<br>(12.9) | 38.8<br>(14.5) | - | 15/15 | 18/13 | - | 0 |
| (Caseras et al., 2013) | - | 31 | 19 | - | 40.5<br>(8.1) | 42.3<br>(6.0) | - | 11/20 | 7/13 | - | 428 |
| (Arrondo et al., 2015) | 22 | 24 | 21 | 32.7<br>(7.6) | 33.1<br>(9.2) | 34.3<br>(10.1) | 19/3 | 17/7 | 17/4 | 401 (91) | 401<br>(91) |
| (Bjork et al., 2004) | - | - | 24 | - | - | 23.8<br>(2.0) | - | - | 6/6 | - | - |
| (Abler et al., 2008) | 12 | - | 12 | 36.7<br>(7.8) | 33.9<br>(11.2) | 36.2<br>(11.2) | 5/7 | 7/5 | 7/5 | 595<br>(357) | 375<br>(397) |
| (MacDonald et al., 2005) | 18 | - | 28 | 27.5<br>(10.2) | - | 25.4<br>(7.5) | 13/5 | - | 18/10 | 0 | - |
| (MacDonald III & Carter, 2003) | 17 | - | 17 | 34.2<br>(7.7) | - | 33.5<br>(5.8) | 12/5 | - | 12/5 | ? | - |
| (Holmes et al., 2005) | 7 | 10 | 9 | 39.0<br>(6.9) | 32.0<br>(9.9) | 34.3<br>(8.1) | 6/1 | 7/3 | 5/4 | ? | ? |
| (Smucny et al., 2018) | 70 | 24 | 53 | 21.0<br>(3.0) | 22.6<br>(2.9) | 21.1<br>(2.8) | 59/11 | 15/9 | 31/22 | 258<br>(214) | 202<br>(109) |
| (Perlstein et al., 2003) | 16 | - | 15 | 36.8<br>(1.9) | - | 36.4<br>(1.8) | 11/5 | - | 9/6 | 175.3<br>(30) | - |
| (Culbreth et al., 2020) | 28 | - | 30 | 37.2<br>(12.3) | - | 35.2<br>(10.6) | 20/8 | - | 23/7 | 311.8<br>(151.5) | - |
| (Huang et al., 2016) | 23 | - | 23 | 34.5<br>(8.2) | - | 34.9<br>(6.7) | 20/3 | - | 15/8 | 312.5<br>(191.7) | - |
| (Prettyman et al., 2021) | 21 | - | 23 | 36.6<br>(7.7) | - | 32.2<br>(10.3) | 12/9 | - | 13/10 | 421<br>(270) | - |
| (Pretus et al., 2021) | 28 | - | 20 | 37.2<br>(8.6) | - | 31.4<br>(8.6) | 18/10 | - | 10/10 | 257<br>(245) | - |
| (Tikász et al., 2019) | 47 | - | 23 | 34.4<br>(9.6) | - | 31.9<br>(8.2) | 47/0 | - | 23/0 | 14.6<br>(8.7) | - |
| (Wolf et al., 2014) | 41 | - | 37 | 41.7<br>(10.7) | - | 39.2<br>(12.1) | 22/19 | - | 18/19 | 493<br>(347) | - |
| (Bi et al., 2022) | - | 33 | 30 | - | 20.7<br>(2.2) | 20.2<br>(1.7) | - | 10/23 | 13/17 | - | 0 |
| (Yang et al., 2016) | - | 25 | 25 | - | 28.9<br>(7.0) | 28.3<br>(7.8) | - | 12/13 | 10/15 | - | 0 |

The detailed information about cluster size and peak ALE maxima and locations are show in Table 2

**Table 2:**
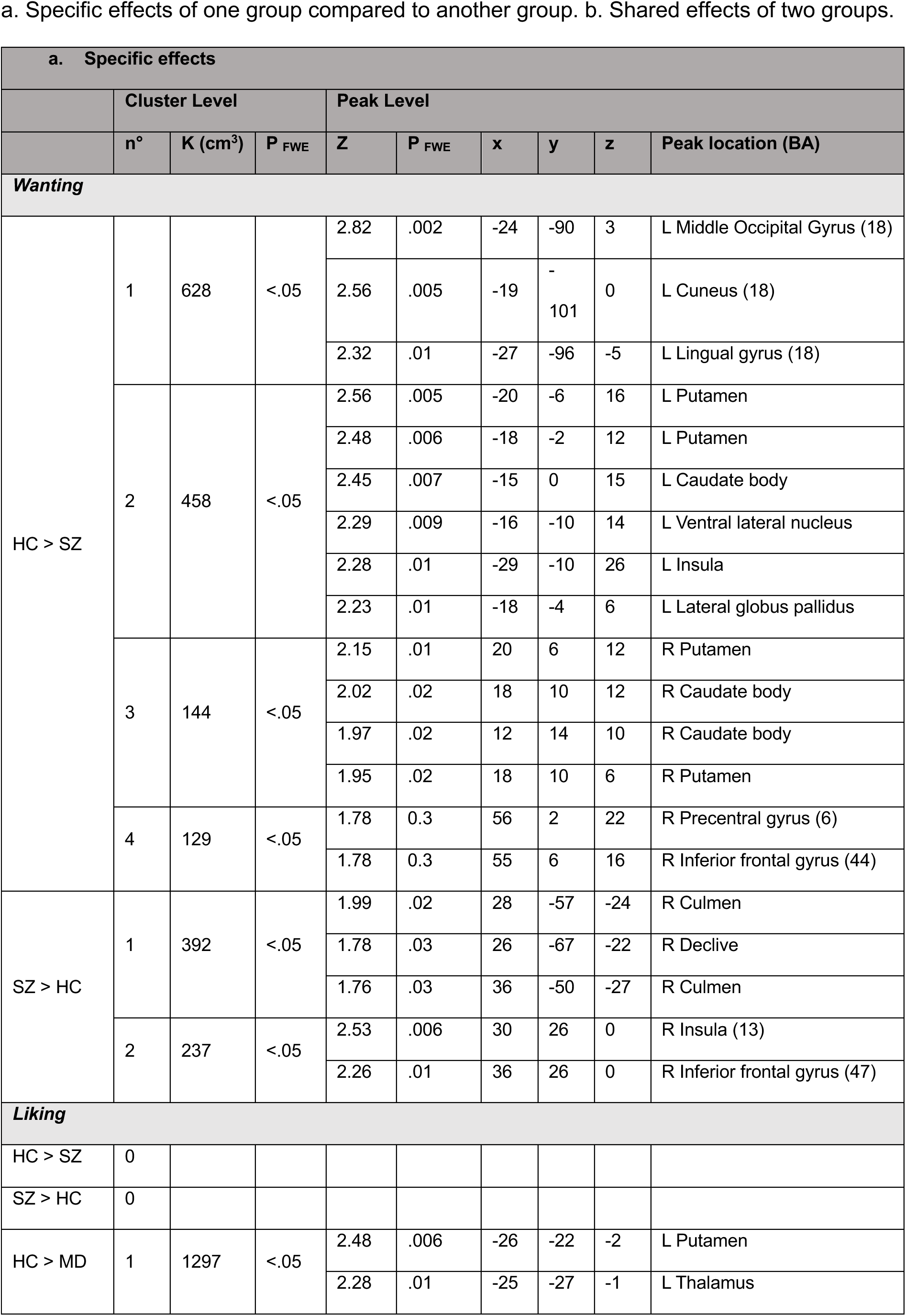
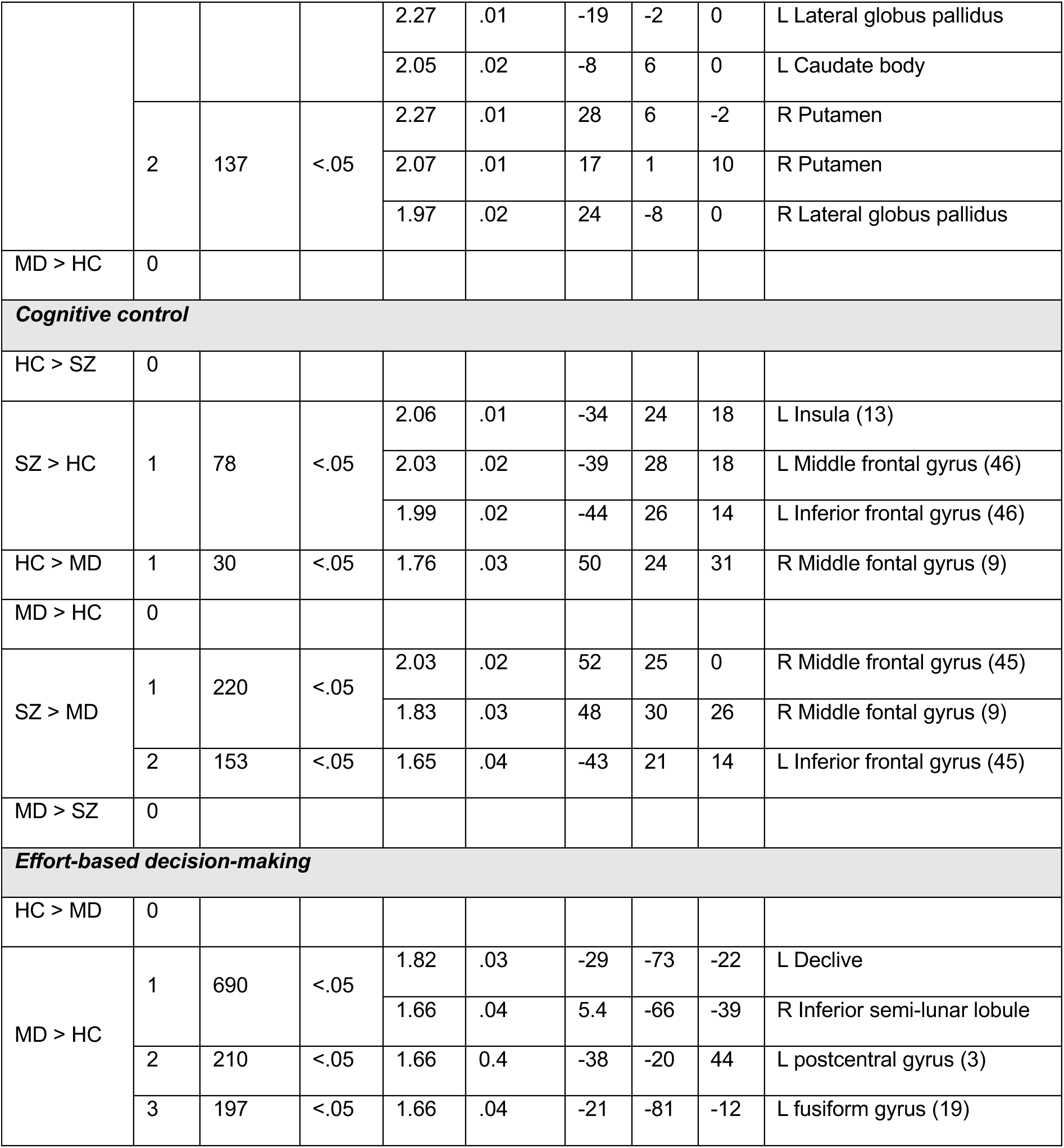
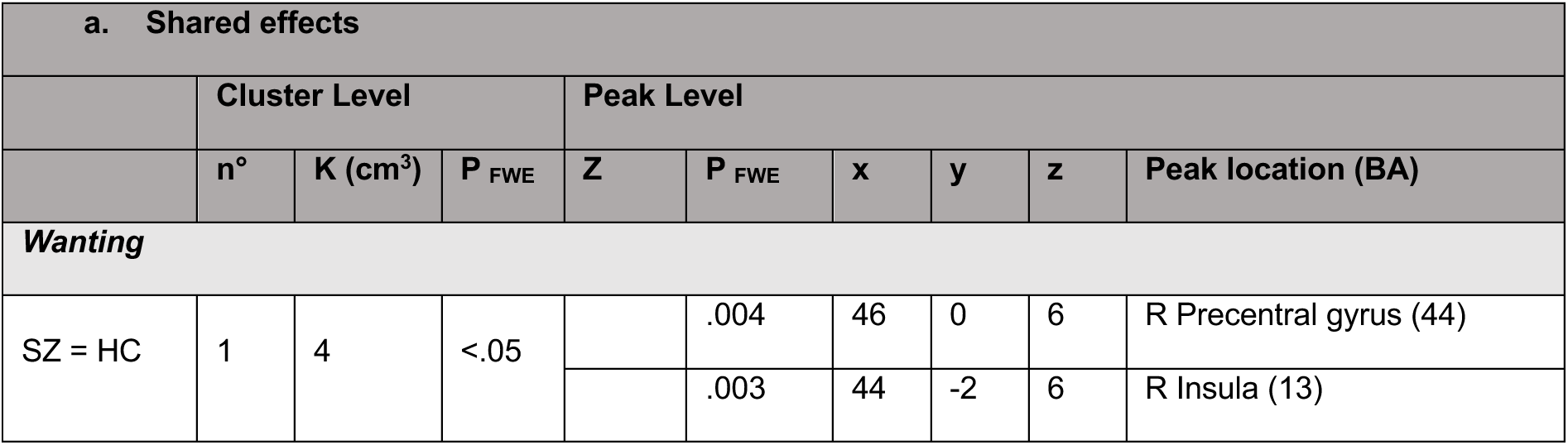
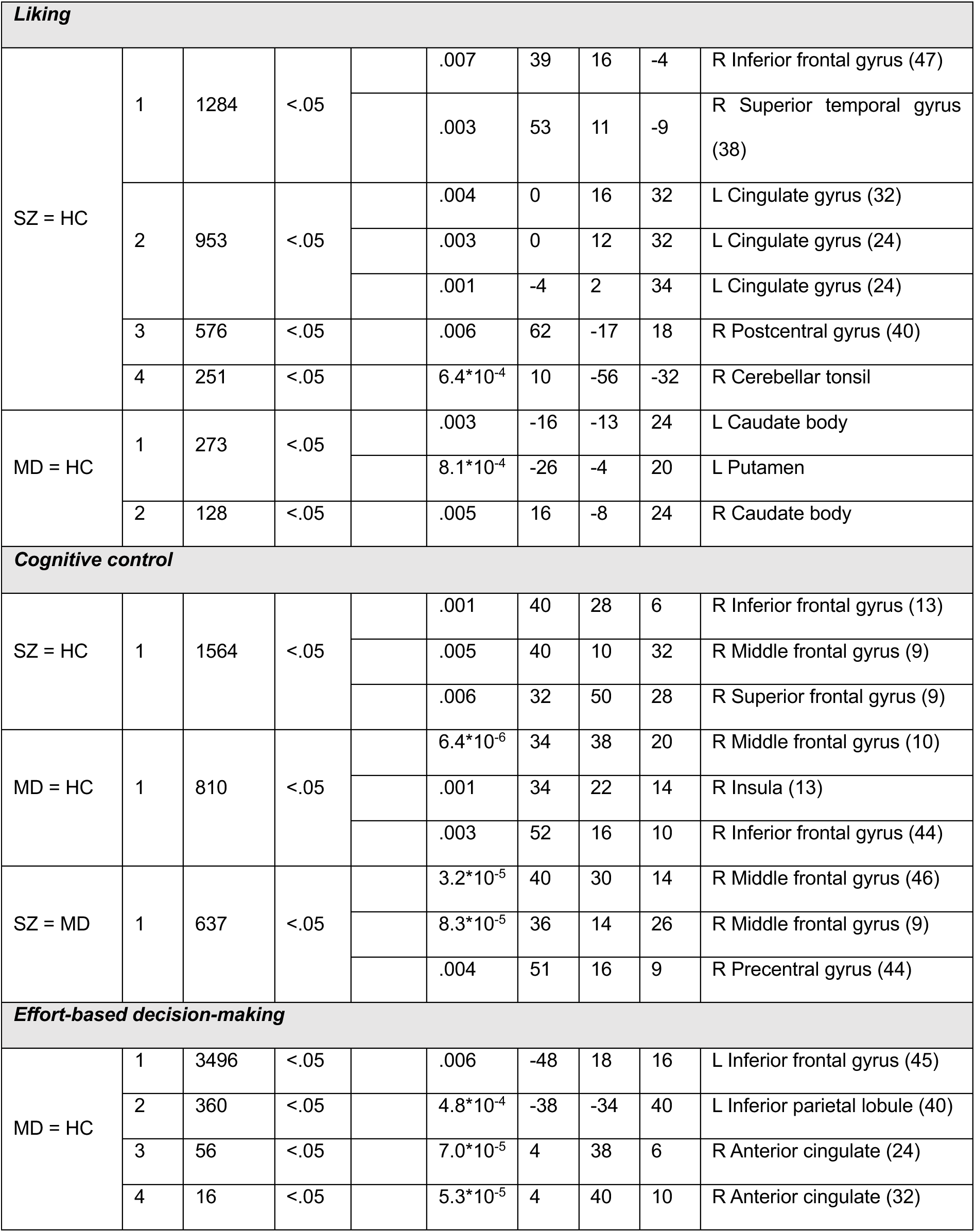
Imaging results. Talairach coordinates x, y, z in mm; BA: Brodmann area. a. Specific effects of one group compared to another group. b. Shared effects of two groups.

### 1. Motivational task

#### 1.1. Wanting (reward/loss > neutral cues)

A total of 15 experiments with 289 HC reporting 170 foci, 4 experiments with 99 MD patients reporting 75 foci, and 11 experiments with 227 SZ patients reporting 94 foci were included in the analyses for wanting.

#### 1.1.1. Within-group analyses

For wanting, ALE analysis revealed, in MD, that no brain regions are consistently found to present a modulation of likelihood of activation after a reward (or loss) cue compared to a neutral cue.

ALE analysis revealed that SZ patients have several regions that present increased activity after a reward (or loss) cue than a neutral cue in right frontal lobe, such as inferior and middle frontal gyrus and insula, as well as in anterior and posterior lobe of the cerebellum, including culmen, declive and uvula.

ALE analysis revealed that HC have increased activity after a reward (or loss) cue compared to a neutral cue that is more frequently reported in a set of subcortical areas, including bilateral putamen, bilateral lateral globus pallidus, bilateral caudate body, precentral gyrus, insula, as well as occipital lobe and right precuneus.

#### 1.1.2. Mood disorders vs. healthy controls

Impossible to do because of the lack of activation in the MD group.

#### 1.1.3. Schizophrenia vs. healthy controls

During reward (or loss) anticipation compared to neutral cue, ALE analysis revealed that, compared to SZ, HC showed increased activity in a set of subcortical areas, including bilateral putamen, left lateral globus pallidus, bilateral caudate body, left ventral anterior and lateral nucleus, right amygdala, right precentral gyrus, as well as left occipital lobe. Conversely, SZ patients present greater activation HC on the right anterior lobe of the cerebellum, including culmen, declive and uvula, as well as the right insula (see Figure 3.a).

**Figure 3:**
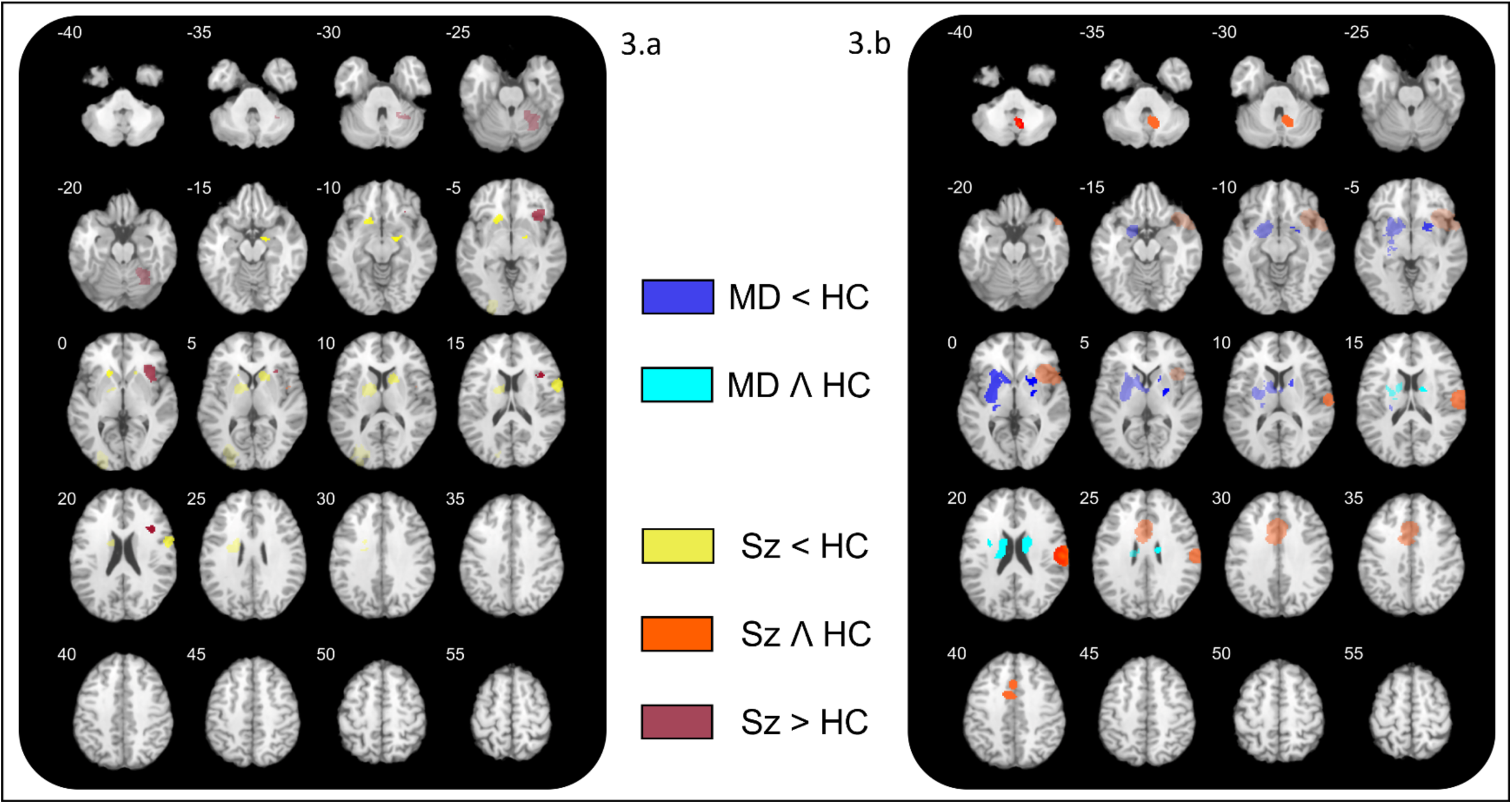
Contrasts maps representing face to face comparison and co-activations. (conjunctions [Λ]) **between patients** (i.e. schizophrenia [Sz] and mood disorders [MD]) **and healthy controls** (HC) **of hyperactivations observed with the Monetary Incentive Delay task.** Numbers correspond to the positioning of the slice in the Z-axis, in mm. ALE meta-analysis (subtraction analysis), p<.05 FWE at the cluster level. 3.a: in “wanting” paradigm; 3.b: in “liking” paradigm.

### 1.2. Liking (reward/loss > neutral feedback)

A total of 11 experiments with 230 HC reporting 120 foci, 4 experiments with 99 MD patients reporting 10 foci, and 7 experiments with 170 SZ patients reporting 9 foci were included in the analyses for liking.

#### 1.2.1. Within-group analyses

For liking, ALE analysis revealed, after reward (or loss) feedback compared to neutral feedback, that MD patients present increased activity of a broad cortical-subcortical network, including left posterior cingulate, left culmen and precuneus, bilateral caudate body, left putamen and bilateral ventral lateral nucleus.

ALE analysis revealed, after reward (or loss) feedback compared to neutral feedback, that SZ patients present increased activity in bilateral frontotemporal gyrus, such as insula, superior temporal gyrus, inferior frontal gyrus, (anterior) cingulate gyrus, as well as in the right posterior-anterior lobe, including uvula, declive and culmen.

ALE analysis revealed, in HC, increased activity after reward (or loss) feedback compared to neutral feedback are consistently found in a bilateral cortical-subcortical network, including putamen, caudate (head and body), lateral globus pallidus, amygdala.

#### 1.2.2. Mood disorders vs. healthy controls

During the reward (or loss) > neutral feedback contrast, HC compared to MD showed greater activation in the cortical-subcortical network, including bilateral putamen, bilateral lateral globus pallidus, left caudate (head and body), left amygdala (see Figure 3.b).

#### 1.2.3. Schizophrenia vs. healthy controls

During the reward (or loss) > neutral feedback contrast, ALE analysis revealed that activations are common to SZ patients and HC (see Figure 3.b).

### 2. Cognitive control tasks (B > A cues)

Because of the lack of articles studying the reactive mode, we focus on the proactive cognitive control, with the anticipation of contextual cues (B vs A cues).

A total of 5 experiments with 122 HC reporting 24 foci, 2 experiments with 34 MD patients reporting 5 foci, and 5 experiments with 128 SZ patients reporting 27 foci were included in the analyses.

#### 2.1. Within-group analyses

ALE analysis revealed that MD patients present an increased activity after a B cue than an A cue that is more frequently reported in several regions in bilateral frontal gyrus, such as right inferior frontal gyrus, left middle / medial frontal gyrus, and anterior cingulate, as well as in the left parietal lobe, including precuneus.

ALE analysis revealed that SZ patients have more consistently an increased activity after a B cue than an A cue in several regions in bilateral frontal lobe, such as middle and inferior frontal gyrus.

ALE analysis revealed that HC present increased activity after a B cue than an A cue in several regions in right frontal lobe, such as middle and inferior frontal gyrus.

#### 2.2. Mood disorders vs. healthy controls

During the B cue > A cue contrast, ALE analysis revealed that HC showed present a greater activation in right middle frontal gyrus compared to MD (see Figure 4).

**Figure 4:**
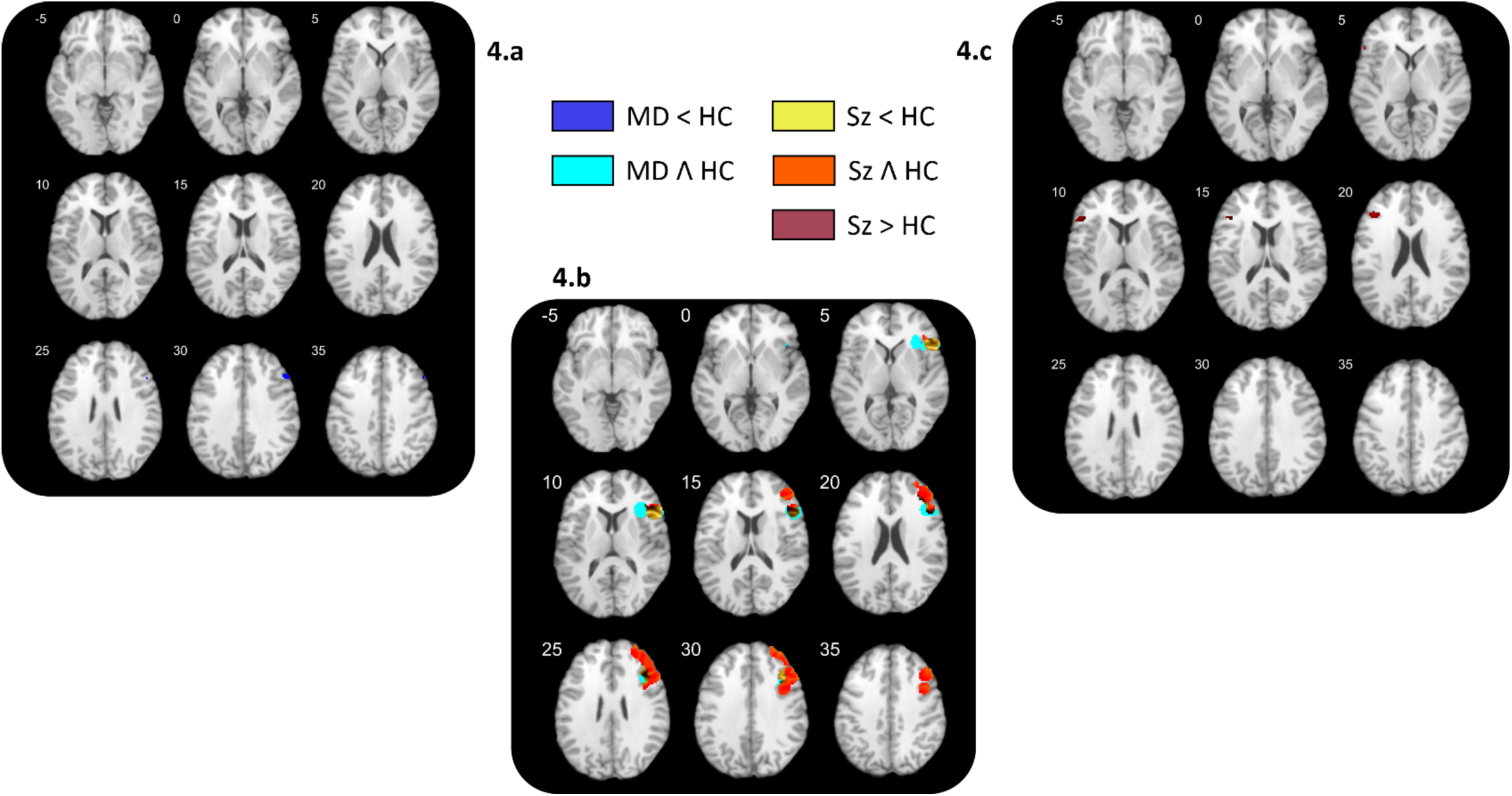
Contrasts maps representing face to face comparison and co-activations. (conjunctions [Λ]) **between patients** (i.e. schizophrenia [Sz] and mood disorders [MD]) **and healthy controls** (HC) **of hyperactivations observed with the AX-Continuous Performance Task (proactive control).** Numbers correspond to the positioning of the slice in the Z-axis, in mm. ALE meta-analysis (subtraction analysis), p<.05 FWE at the cluster level. 4. a: when activation in HC are superior to those in other conditions 4.b: common activations between HC and patients 4.c: when activation in HC are inferior to those in other conditions

#### 2.3. Schizophrenia vs. healthy controls

During the B cue > A cue contrast, ALE analysis revealed that SZ patients compared to HC showed increased activation in left inferior and middle frontal gyrus (see Figure 4).

### 3. Effort-based decision-making tasks (high effort > low effort)

A total of 8 experiments with 211 HC reporting 28 foci, 2 experiments with 58 MD patients reporting 11 foci, and 6 experiments with 188 SZ patients reporting 22 foci were included in the analyses.

#### 3.1. Within-group analyses

ALE analysis revealed that MD patients present an increased activity after a high-effort cue than a low-effort cue, which is more frequently reported in a broad cortical-subcortical network, including left middle / inferior frontal gyrus, right medial frontal gyrus, left precentral gyrus, left postcentral gyrus, as well as left declive and right inferior semi-lunar lobule.

ALE analysis revealed, in SZ, that no brain regions were consistently found to present a modulation of likelihood of activation after a high-effort cue compared to a low-effort cue.

ALE analysis revealed that HC present increased activity after a high-effort cue than a low-effort cue in several regions in a broad cortical-subcortical network, such as left medial frontal gyrus, bilateral anterior cingulate, bilateral inferior frontal gyrus, right caudate head, bilateral insula and left supramarginal gyrus.

#### 3.2. Mood disorders vs. healthy controls

During the high-effort cue > low-effort cue contrast, ALE analysis revealed that MD showed presented a greater activation in the left posterior cerebellum (declive), left postcentral gyrus, left fusiform gyrus, and right posterior cerebellum (inferior semi-lunar lobule) compared to HC (see Figure 5).

**Figure 5:**
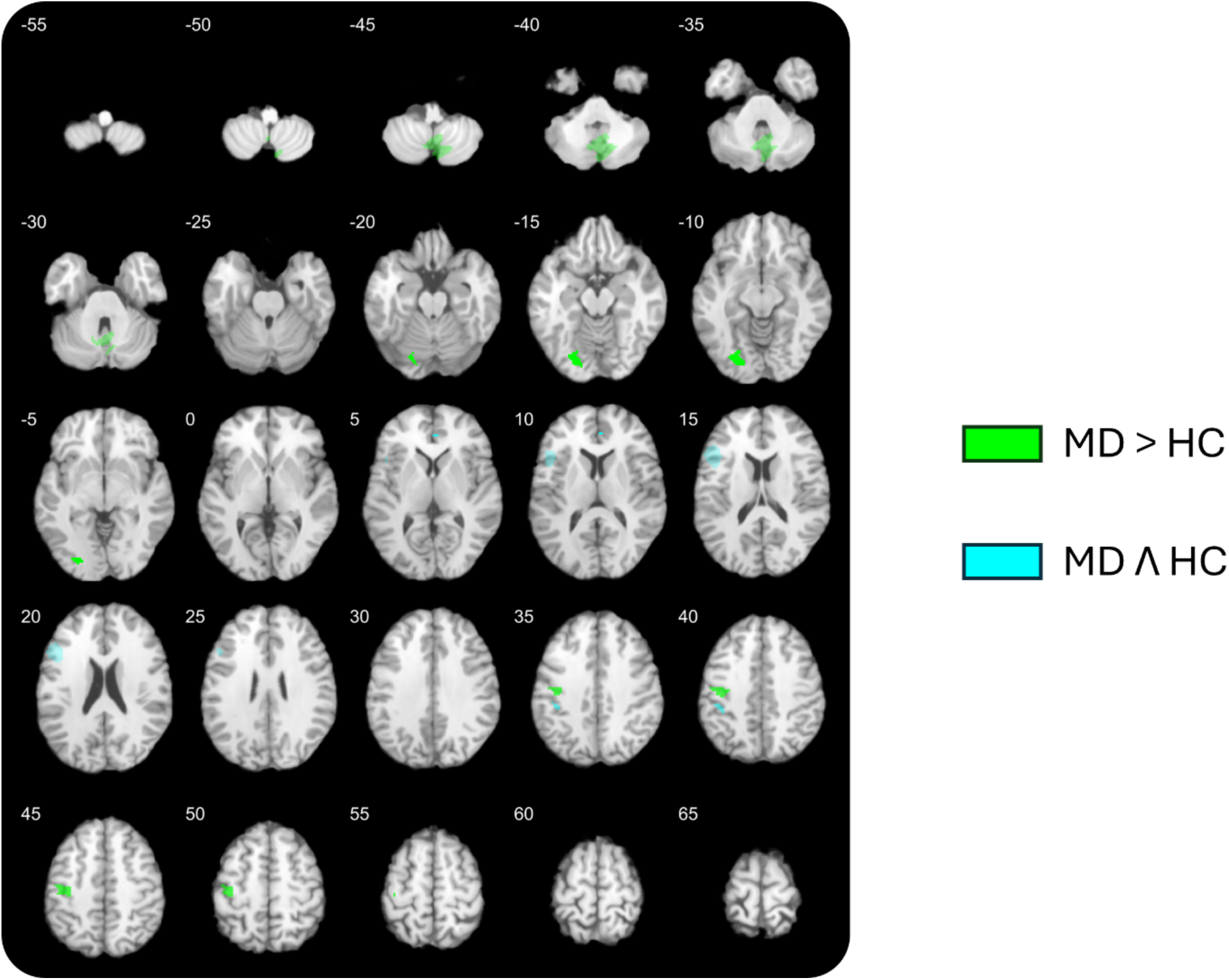
Contrasts maps representing face to face comparison and co-activations. (conjunctions [Λ]) **between patients** (i.e. mood disorders [MD]) **and healthy controls** (HC) **of hyperactivations observed with the effort-based decision-making tasks.** Numbers correspond to the positioning of the slice in the Z-axis, in mm. ALE meta-analysis (subtraction analysis), p<.05 FWE at the cluster level.

#### 3.3. Schizophrenia vs. healthy controls

Impossible to do because of the lack of activation in the SZ group.

## Discussion

This ALE meta-analysis of neuroimaging studies of motivational, cognitive control, and effort-based decision-making networks in SZ and MD has highlighted that patients suffering from psychiatric disorders, in comparison to HC, present specific patterns of activation. Moreover, these patterns are distinct for SZ and MD and thus could suggest, based on the multidimensional neurological model of apathy, the existence of different dominant forms of apathy.

Concerning SZ, during the MID, SZ patients present the same activation as HC during the feedback phase, in accordance with the literature about a preserved liking in SZ [113,114]. However, during the cue phase, compared to HC, SZ patients present more frequent hypoactivation of the motivational network (bilateral putamen, left lateral globus pallidus, bilateral caudate body, left ventral anterior and lateral nucleus, right amygdala). This result, suggesting impaired wanting in SZ [115], had been already linked to apathy and anhedonia [95,100,116,117]. However, interestingly, during the cue phase, SZ patients also present more consistent hyperactivation than HC of the right cerebellar vermis (culmen, declive and uvula). The cerebellar vermis in the cerebellum is a well-known structure for the regulation of emotions and affects [118,119]. Theta burst stimulation of this structure in SZ has previously been shown to offer the potential to modulate emotions and affects in SZ patients [120]. Since the cerebellum is highly connected to the motivational network and to the dorsolateral and ventral PFC [121–123], the hyperactivation of the cerebellar vermis in SZ could thus be a sort of compensation for the hypofunctionality of the wanting network. The hyperactivation of the cerebellar vermis highlighted specifically in SZ in the present study, and which has previously been correlated to the severity of apathy and emotional blunting in this pathology [124], could therefore suggest a dominant emotional-affective form of apathy in SZ. Results obtained with AX-CPT strengthen this hypothesis in a way that excludes the cognitive one. Indeed, in comparison to HC, SZ patients present a more frequent hyperactivation of the frontal cognitive structures, especially the bilateral frontal gyrus, known to support the execution of attentional and cognitive control [125,126]. This hyperfrontality in some regions of the PFC in SZ had previously been highlighted to reflect a shift in strategy in an attempt to compensate for the proactive difficulty often shown by SZ patients [105,127–130]. Accordingly, an ALE meta-analysis of executive functioning in SZ confirmed the hyperactivation of the midline areas as a compensatory response for the executive difficulties and/or as an alternate attentional strategy to support task performance [131]. Using such strategies do not fit with the hypothesis of a dominant cognitive form of apathy in SZ, with cognitive apathy being precisely associated with a reduced functioning of the executive and attentional networks [16,17,132]. As suggested in the only study that has investigated the three forms of apathy in SZ, via a survey and neuropsychological tests, SZ patients with emotional apathy could actually be even more prone to using this compensatory strategy by allocating more attentional resources to non-emotional stimuli [133]. In the effort-based decision-making tasks, SZ patients showed a lack of specific activation between low and high effort choices, suggesting an impaired decision-making processing. This result has been correlated to apathy in several studies, for both cognitive effort [42–44,54,55,60], and physical effort [40,41,44,48,50,53,56–59]. These results could suggest a coexistence of both emotional and initiative apathy in SZ. However, the behavioral studies using effort-based decision-making tasks showed that the more apathetic the SZ patients are, the less effort they expend only for higher rewards. Therefore, the results suggest that SZ impairments during decision-making linked to apathy are possibly due to an impairment of reward rather than effort processing, following a difficulty to translate reward cues into selecting and executing appropriate actions. In favor of this hypothesis, several studies have shown that effort avoidance (cognitive or physical) during effort-based decision-making was not associated with apathy in SZ [45–47,49,51,52]. In conclusion, motivational wanting impairment, a core deficit in SZ, may render everyday tasks abnormally effortful for psychotic patients, inducing a reduction of goal-oriented behaviors, i.e. apathy. This impairment could therefore suggest a dominant emotional-affective form of apathy in at least some subtypes of SZ, even if coexistence of initiative apathy could be possible. However, the preeminence of emotional alterations may be inconstant according to the SZ subtype. For example, progressive periodic catatonia (PCC), a SZ subtype marked by alteration of psychomotricity, was associated in perfusion MRI with hyperactivity in the left premotor cortex that is more associated with alteration in the initiation of the motor-related part of behavioral process [134].

By highlighting impaired functioning of wanting and liking (MID), as well as cognitive control (AX-CPT), the results of the three tasks in the MD patients suggest further a dominant mixed form of apathy in MD: the co-existence of emotional and executive apathy. Indeed, firstly, in the MID, MD patients show a total lack of activation during the cue phase, suggesting an impaired wanting. This result had never been correlated to apathy, but could rather reflect a specific marker of the MD pathology [98,100,117]. At the feedback reception, in comparison to HC, they present more frequent hypoactivation, even if they recruit larger structures of the motivational network (bilateral putamen, bilateral lateral globus pallidus, left caudate, left amygdala). This hypoactivation of the motivational network, suggesting a liking deficit, had already been linked to apathy in MD [98]. Then, regarding the results highlighted in the present meta-analysis with the AX-CT, compared to HC, MD patients present more consistent hypoactivation of the PFC, as well as the right middle frontal gyrus and left inferior frontal gyrus, two structures supporting cognitive control. In the literature, the reduced connectivity of these PFC regions has previously been shown in MD and correlated to the severity of the MD [135]. Such cognitive disruptions in MD have been previously related to inhibition and shifting impairments [38,136]. Interestingly, and despite the vast heterogeneity in MD, the reduced activation of the PFC during cognitive control tasks has been correlated to a specific anhedonic phenotype of depression, confirming the combination of mixed impairments, both in the cognitive control and motivation domains [106,137]. Finally, concerning effort-based decision-making tasks, MD patients present the same activation as HC of the decision-making cortical-subcortical network, including anterior cingulate cortex, suggesting a preservation of decision-making processing in MD. However, patients with MD also hyperactivate the left declive during high-effort choices compared to HC. This posterior part of the left cerebellar vermis has been identified in the affective-limbic network, mainly implicated in social cognition and emotional regulation [138]. In two studies including a meta-analysis, an increased activation of this structure was not linked to apathy, but rather to the severity of depressive symptoms in MD patients with and without medication, suggesting that this hyperactivation is a potential marker of depression but irrespective of specific processes such as those related to apathy [139,140]. Does this mean that these results correspond to pathophysiological or etiological mechanisms common to MD and SZ, insofar as they are marked by apathy? We believe that we can make no such claim, for several reasons. Firstly, because these observations come from heterogeneous populations, it is now well accepted that there are subtypes of SZ, as well as in MD, and that many of them are accompanied by apathetic symptoms. This heterogeneity implies substantial inter-individual variability in functional imaging studies [141,142]. Secondly, while experimental conditions may have reduced this heterogeneity, they also focused on certain neurological systems more involved in the tasks, thus underestimating lesser-involved dysfunctions further up the etiological causal chain. It is quite possible that we have identified downstream activity modifications in the processes affected by the various forms of apathy and that these modifications are common to the different pathologies by the equifinality effect [12]. A new analysis considering the differences between clinical phenotypes within each diagnosis and exploring the different networks that may influence the regions we have identified, including under ecological conditions, would complete our description of the pathophysiology of apathy during psychiatric diseases.

This study has some limitations, the main one being the use of one database, the relatively small number of studies included and their own small population sizes. Some limits are inherent to the literature to date, with a lack of results for some contrasts and paradigms and poor control on the treatment, age, and severity of the disorders. Moreover, psychiatric diagnoses are based on DSM classification: if populations are assumed to be comparable, they are also likely to be heterogeneous within the groups themselves and according to their pathophysiological processes. This limitation may be particularly true for SZ, which constitutes a particularly heterogeneous group with marked differences in clinical presentations, underlying mechanisms, and ease of inclusion in imaging studies. Finally, this study is a coordinate-based meta-analysis of studies that lack power, which can create some false negatives, especially in the case of heterogeneous groups [73]. However, the realization of image-based meta-analyses required unavailable, highly detailed data. If they were available, their analysis could also offer possibilities for phenotyping according to this three-dimensional model of apathy.

Relying on an ALE meta-analysis of neuroimaging studies that have investigated motivational or cognitive control or effort-based decision-making processing in SZ or MD, we highlighted specific impairments in each of these two pathologies. The discussion of these specificities, regarding the known neural substrates of multidimensional apathy, allowed us to suggest a possibly dominant emotional-affective form of apathy in SZ and the coexistence of emotional-affective and executive forms of apathy in MD, which could support the development of new therapeutic strategies in precision psychiatry.

## Acknowledgements

This research was supported by the French Ministry of Higher Education and Research grant (for the PhD of G. L-B). G. L-B and A. B designed the study. G. L-B and L. C. D-J identified the papers and realized the analyzes. G. L-B wrote the first draft of the manuscript. All the authors were involved in the writing of this draft.

